# The IGVF Catalog Knowledge Graph

**DOI:** 10.64898/2026.11.20.768224

**Authors:** Shengcheng Dong, Ian Whaling, Otto Aleksi Jolanki, Jennifer Jou, Zhewei Shen, Khine Zin Lin, Pedro Assis, Idan Gabdank, Keenan Graham, Meenakshi Kagda, Mingjie Li, Forrest Tanaka, Jesse M. Engreitz, Shane Liu, Daofeng Li, Ting Wang, Michael I Love, Lars Steinmetz, Benjamin C Hitz

**Affiliations:** Department of Genetics, School of Medicine, Stanford University, Stanford, CA 94305; Department of Biomedical Data Science, School of Medicine, Stanford University, Stanford, CA 94305; Department of Genetics, The Edison Family Center for Genome Sciences and Systems Biology, Washington University School of Medicine, St. Louis, MO 63110, USA; Department of Genetics, University of North Carolina at Chapel Hill, Chapel Hill, NC, 27599, USA; Department of Biostatistics, University of North Carolina at Chapel Hill, Chapel Hill, NC, 27599, USA

## Abstract

The Impact of Genomic Variation on Function (IGVF) consortium is systematically characterizing how genomic variation influences regulatory elements, gene expression, and human phenotypes through thousands of experimental mapping and perturbation datasets alongside predictive models. To synthesize these diverse results into quantitative relationships and situate them within established biological knowledge, we developed the IGVF Catalog, a large-scale knowledge graph. Implemented in ArangoDB using schema-validated JSON records. Version 1.3 encompasses nearly 3 billion nodes across 14 entity types (including variants, coding variants, genomic elements, genes, transcripts, proteins, pathways, and ontology graphs for biosamples and phenotypes) connected by more than 8.6 billion edges across 31 relationship types. Edges encode statistical significance, effect sizes, source provenance, and biological context via ontology-linked hyperedges. The Catalog integrates 657 IGVF datasets spanning non-coding reporter and CRISPR perturbation screens (MPRA, STARR-seq, Perturb-seq), coding variant assays (VAMP-seq, SGE, MAVE, DUAL-IPA), and predictive models (scE2G, SEMVAR, cV2F, TLAND, and BlueSTARR), unified with extensive public context including GENCODE annotations, FAVOR and dbSNP variants, TopLD linkage disequilibrium, GWAS, QTLs, and protein interactions. Users can interrogate the Catalog through a web portal (https://catalog.igvf.org), Python and R clients, an OpenAPI-documented API featuring 86 endpoints supporting identifier and genomic-region queries, and a Model Context Protocol (MCP) server enabling agent-assisted querying with large language models. We demonstrate the Catalog’s utility for mechanistic interpretation across three case studies: resolving a FOXA1-dependent enhancer variant (rs11257655) regulating *CAMK1D* at a type 2 diabetes locus, distinguishing coding (*GCKR* rs1260326) and regulatory (*HNF1A* rs1169289) mechanisms for liver traits, and linking vascular CRISPRi-perturbed elements to *CDKN2A*/*CDKN2B* and *NOS3* in coronary artery disease. Updated quarterly with open data and software, the IGVF Catalog provides a scalable resource for variant-to-function discovery.

## Introduction

The Impact of Genomic Variation on Function consortium was established to continue the study of the human genome at scale [1]. Over 100 labs across 25 awards have produced over 2500 experimental datasets across dozens of single cell mapping and perturbation assays in hundreds of cellular systems. In addition, predictive modeling groups have supplied broad and deep predictions of variant and element effects. The IGVF Catalog was created to distill this vast body of work into quantitative relationships between variants, elements, genes, and phenotypes, and to put these data in the context of public biological and genomic databases and results. To this end, we have created a large knowledge graph representing 14 node types and 31 edge types between these nodes. All records (nodes and edges) are represented in the database as JSON objects. A “knowledge graph” is a specific type of graph database where the entities have a set of properties, the properties in the JSON objects, associated with them. For example, a gene has a start and stop coordinate on the reference genome (GRCh38 for human, GRCm39 for mouse), a strand, a name (HGNC) [2], various identifiers (the Ensembl ID is used as the primary key [3]), a gene type (e.g. protein coding), and several aliases. Nodes represent genomic entities (genes, variants, genomic elements), derived biological entities (coding variants, gene structure, transcripts, proteins, complexes, pathways), and a few other items (studies, donors, drugs). Phenotypes (including diseases) and biosamples (tissues, cell lines, cell types, etc.) are represented by full ontology graphs (directed acyclic graphs; DAGs). The ontologies currently loaded in the Catalog (both terms as nodes and relationships as edges) are listed in **Table 1**. Edges represent relationships between nodes. As an example, a (protein coding) gene has edges to its transcripts and its transcripts have edges to a single protein. Genes also have edges to ontology terms; specifically biological function, process, and cellular component ontology terms from the Gene Ontology (GO) [4]. Other edges represent experimental observations or predictions; for example an expression quantitative trait locus (eQTL) is represented by an edge between a variant and and a gene, with that edge’s properties representing the statistical significance and effect size of the relationship, along with the source of this information (either an external database or an IGVF product). Within the catalog there are many examples of “hyperedges”, that is a set of edges (usually 2-3) with a set of nodes (usually 3-4). In the eQTL example above, that statistical observation of the relationship between variant and gene takes place in a particular biological context (a tissue or cell type, for example); this meta-edge is captured as a property of the edge that acts as a pointer to an ontology term (biosample) node. Using this type of logic, we have constructed a discrete model for nearly all of the data produced by the IGVF consortium as well as put it into context with public genomics data. In total, the knowledge graph (version 1.3) contains nearly 3 billion nodes and more than 8.6 billion edges. A somewhat simplified version of how various types of experimental and prediction data are modeled in the knowledge graph is shown in **Figure 1**. All nodes have _key, name, source, and source_url properties. Nodes derived from IGVF data (usually genomic elements but sometimes variants) have a files_fileset property and a method property, indicating what the specific source file was, and what method (experimental assay type or predictive software) was used to define it. Node names are usually database identifiers or other common unique labels. All edges have _key, _from, _to, source, source_url, name and inverse_name. _from and _to are special properties of an edge that connect the graph (typically the _key property of nodes). This is what makes the document-style database of JSON objects into a *graph*. This doesn’t always imply directionality. The name field of an edge is meant to be read as (_from.name) - (name) - (_to.name). For example, an edge in the variants_proteins collection, derived from the SEMVAR predictions of allele specific binding of a TF [5] has the following properties:

**Table 1.**
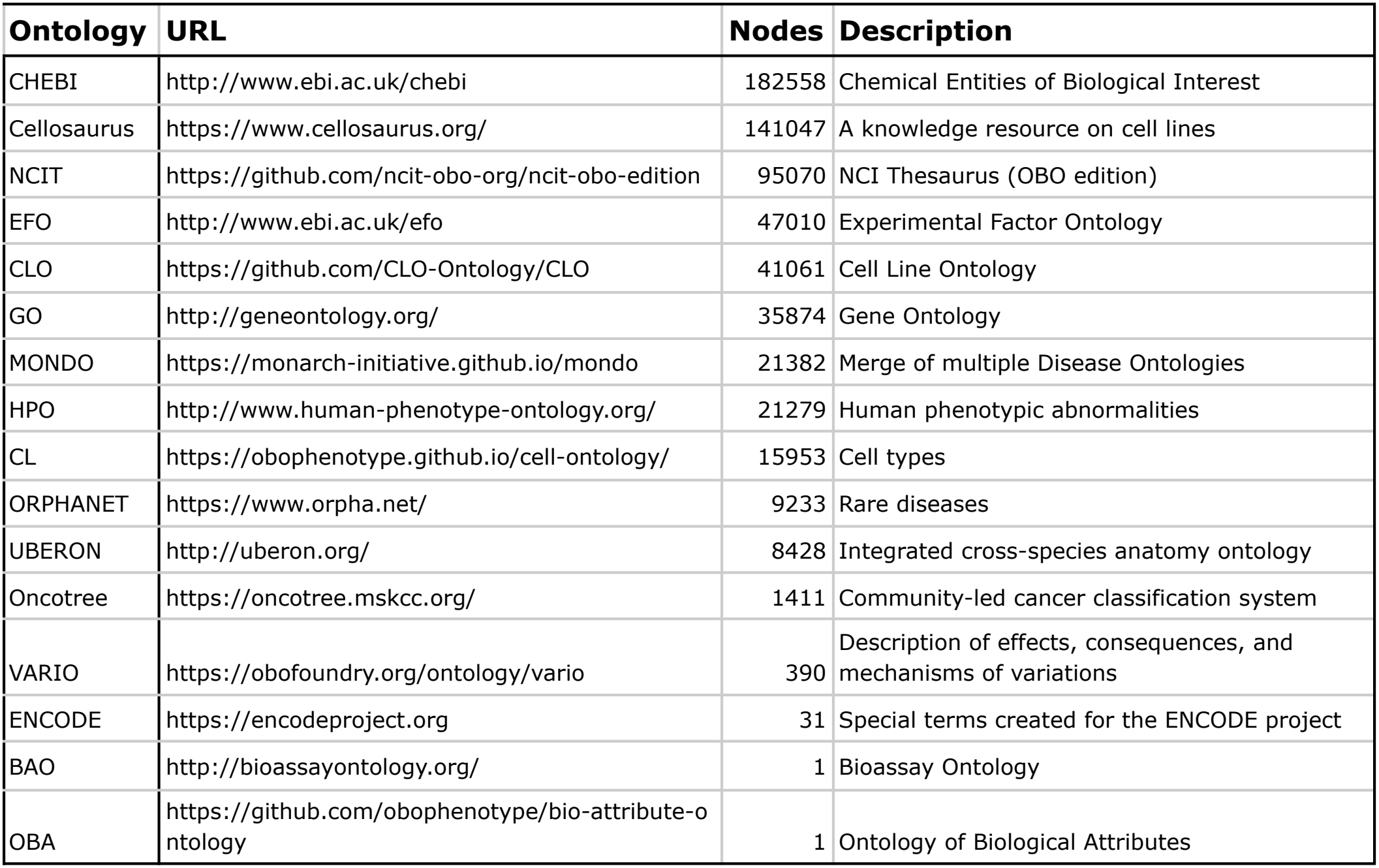
List of Ontologies in the IGVF Catalog. This table shows each ontology collection’s source url, the number of term nodes loaded in the IGVF Catalog.

| **Ontology** | **URL** | **Nodes** | **Description** |
|---|---|---|---|
| CHEBI | http://www.ebi.ac.uk/chebi | 182558 | Chemical Entities of Biological Interest |
| Cellosaurus | https://www.cellosaurus.org/ | 141047 | A knowledge resource on cell lines |
| NCIT | https://github.com/ncit-obo-org/ncit-obo-edition | 95070 | NCI Thesaurus (OBO edition) |
| EFO | http://www.ebi.ac.uk/efo | 47010 | Experimental Factor Ontology |
| CLO | https://github.com/CLO-Ontology/CLO | 41061 | Cell Line Ontology |
| GO | http://geneontology.org/ | 35874 | Gene Ontology |
| MONDO | https://monarch-initiative.github.io/mondo | 21382 | Merge of multiple Disease Ontologies |
| HPO | http://www.human-phenotype-ontology.org/ | 21279 | Human phenotypic abnormalities |
| CL | https://obophenotype.github.io/cell-ontology/ | 15953 | Cell types |
| ORPHANET | https://www.orpha.net/ | 9233 | Rare diseases |
| UBERON | http://uberon.org/ | 8428 | Integrated cross-species anatomy ontology |
| Oncotree | https://oncotree.mskcc.org/ | 1411 | Community-led cancer classification system |
| VARIO | https://obofoundry.org/ontology/vario | 390 | Description of effects, consequences, and mechanisms of variations |
| ENCODE | https://encodeproject.org | 31 | Special terms created for the ENCODE project |
| BAO | http://bioassayontology.org/ | 1 | Bioassay Ontology |
| OBA | https://github.com/obophenotype/bio-attribute-ontology | 1 | Ontology of Biological Attributes |

**Figure 1.**
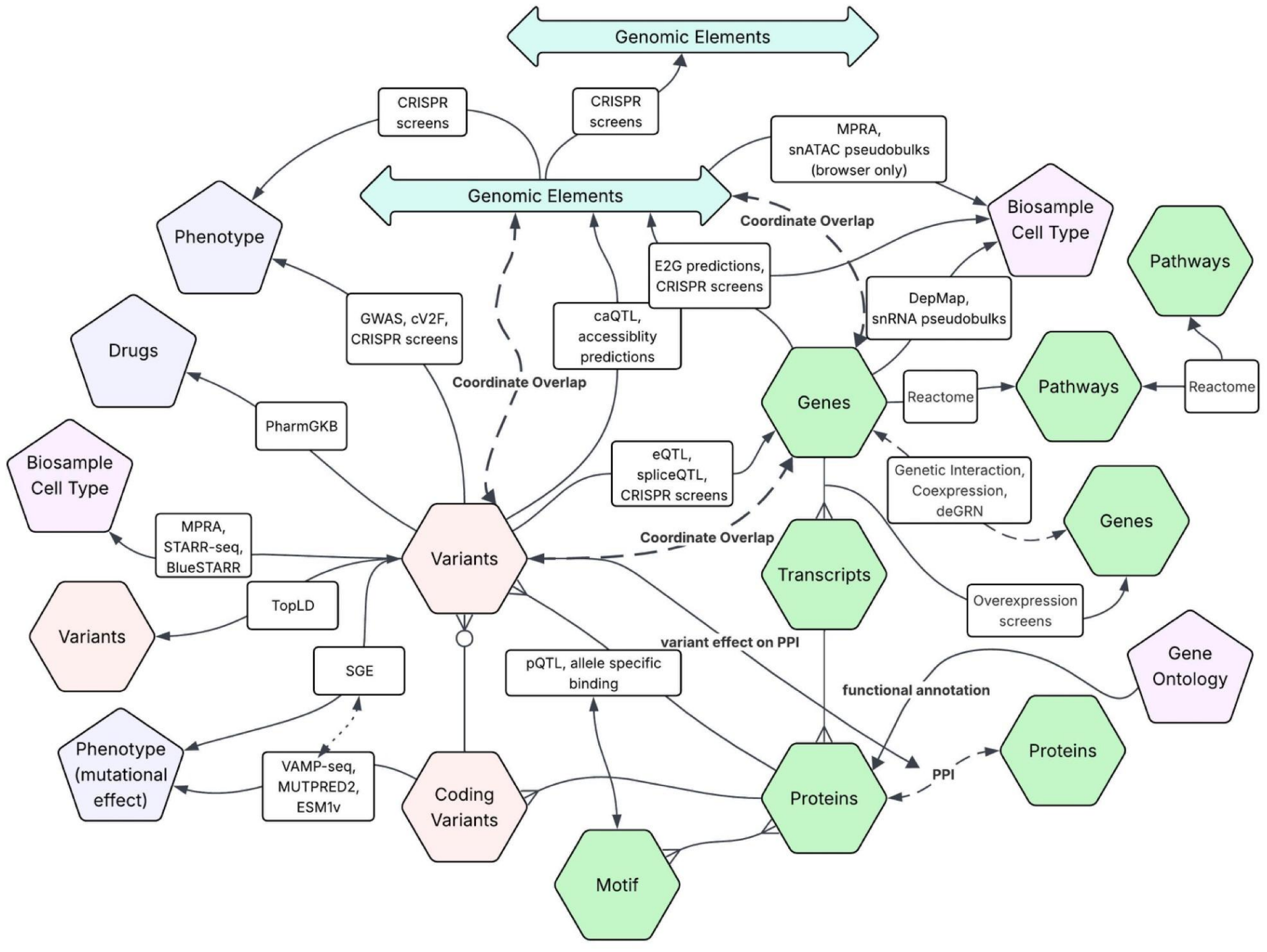
Knowledge Graph relationships. Schematic representation of the knowledge graph. Gene/protein nodes are in green, variants and protein coding variants in light red and ontology terms are in purple. Connections between nodes (edges) are labeled by corresponding *methods* in the database. The large blue arrow represents genomic element nodes, and the dashed connections represent the possibility to query by genomic interval. Gene-gene and protein-protein edges are true edges but are distinguished as edges between nodes of the same type. Pseudobulks from snATAC and are not loaded as true edges, but references to genome browsable files in the files_filesets metadata collection. SGE (saturation genome editing) is a method which assesses the effect of coding mutations, but is performed at the nucleotide level, hence it has been encoded as both a genomic variant and protein coding variant effect. *Not shown*: Gene programs, edges between ontology terms (loaded from OWL DAG files), gene structure, studies (from GWAS), drugs, other ontology terms not representing phenotypes, protein function, or biosamples.

~~~
{
_from: variants/NC_000001.11:1000017:G:A,
_to: proteins/ENSP00000381518, name: “modulates binding of”,
inverse_name: “binding modulated by”,
}
~~~

(ENSP00000381518 is the protein produced by the transcription factor gene MEIS1). This can be “read” as “the G->A variant on Chromosome 1, position 1000017 modulates binding of MEIS1. The inverse name works the same way but reversing the _from, and _to, so this record can also be read “MEIS1 binding modulated by the G->A variant on Chromosome 1, position 1000017”. Most edges also have a class and a method (see API description). The Catalog can also be accessed via a user interface at https://igvf.catalog.org [6].

## Results

### IGVF Data in the Catalog

The primary repository for all IGVF Data (files and associated metadata) is the IGVF Portal (https://data.igvf.org; Jou and Shen *et al*, in preparation). Not every file or data point in the portal is represented in the catalog, but the datasets and files that are are labeled by a “collections” property allowing them to be searched and filtered from the Portal. Version 1.3 contains 657 IGVF datasets (https://data.igvf.org/search/?type=FileSet&status=released&collections=IGVF_catalog_v1.3&award.project=IGVF&award.project=IGVF+affiliate), with roughly one file per dataset loaded. All files loaded in the IGVF Catalog version 1.3, along with their source, producing labs, methods and sample information are listed in **Supplementary Table 1**. With some exceptions, IGVF experiments and predictions are represented by *edges* in the Catalog. IGVF has experiments and predictions assessing non-coding variants, coding variants, and relationships between genomic elements (putative enhancers or promoters) and genes. Most IGVF experimental edges characterizing noncoding regulation are derived from reporter assays, including MPRA and STARR-seq [7,8], and CRISPR screens, such as Perturb-seq [9]. Edges derived from these screens include inferred gene regulatory networks.

Coding variant experiments (and associated predictions), using techniques such as VAMP-seq [10], SGE [11], DUAL-IPA [12], or MAVE [13] assess the effect of coding changes (usually amino acid, but other sites as well in the case of SGE) on protein stability, cell viability, or other readouts. Other predictions in the IGVF corpus include allele-specific effects on transcription factor binding (SEMVAR [5]), predictions of variant effects on phenotypes including gene expression (cV2F, TLAND, BlueSTARR [5,14–16]) and enhancer-gene predictions (scE2G) [17]. The number of edges of each type, from each experiment or prediction is detailed in **Table 2**. Data from the IGVF Data Portal were incorporated into the Catalog using standardized field names for statistical significance, effect sizes, and other scores. Experimental results were generally retained without filtering by statistical significance, whereas predictions and data from public databases were filtered using dataset-specific thresholds to limit database size and support efficient queries.

**Table 2.** List of IGVF datasets in the Knowledge Graph. This table lists the number of records from IGVF experiments and predictions, grouped by collection name, entity measured, edge type and method.

| **BIOLOGICAL RELATIONSHIP** | **EDGE TYPE** | **METHOD** | **CLASS** | **COUNT** |
|---|---|---|---|---|
| coding variant effect | coding variants-phenotypes | VAMP-seq | observed data | 58,212 |
| coding variant effect | coding variants-phenotypes | VAMP-seq (MultiSTEP) | observed data | 135,650 |
| coding variant effect | coding variants-phenotypes | SGE | observed data | 15,143 |
| coding variant effect | coding variants-phenotypes | DUAL-IPA | observed data | 791 |
| coding variant effect | coding variants-phenotypes | Variant painting via fluorescence | observed data | 794 |
| coding variant prediction | coding variants-phenotypes | MutPred2 | prediction | 641,276,378 |
| coding variant prediction | coding variants-phenotypes | ESM-1v | prediction | 454,527,037 |
| ORF overexpression effect | genes/transcripts-genes | MORF screen | observed data | 8,083 |
| element effect on regulatory activity | genomic elements-biosamples | MPRA | observed data | 406,488 |
| element effect on gene expression | genomic elements-genes | CRISPR screen | observed data | 15,171 |
| element effect on gene expression | genomic elements-genes | Perturb-seq | observed data | 58,354,725 |
| element-gene links | genomic elements-genes | scE2G | prediction | 12,416,701 |
| element effect on chromatin accessibility | genomic elements-genomic elements | Multiome Perturb-seq | observed data | 10,935 |
| element effect on phenotype | genomic elements-phenotypes | CRISPR screen | observed data | 1,940 |
| element effect on gene expression (mouse) | genomic elements-genes (mouse) | Perturb-seq | observed data | 257,900 |
| TF motif-protein binding | motifs-proteins | SEMpl | observed data | 820 |
| variant effect on regulatory activity | variants-biosamples | STARR-seq | observed data | 63,940,196 |
| variant effect on regulatory activity | variants-biosamples | MPRA | observed data | 385,440 |
| variant effect on regulatory activity | variants-biosamples | BlueSTARR | prediction | 11,708,699 |
| variant effect on gene expression | variants-genes | CRISPR screen | observed data | 984 |
| variant effect on phenotype | variants-phenotypes | CRISPR screen | observed data | 10,987 |
| variant effect on phenotype | variants-phenotypes | SGE | observed data | 24,391 |
| variant effects | variants-phenotypes | cV2F | prediction | 2,046,197 |
| variant effect on TF binding | variants-proteins | SEMVAR | prediction | 357,795,432 |

### Biological Context

One of the primary motivating design factors for the IGVF Catalog was to put the results and predictions from the consortium into context relative to known biology. To this end we have loaded many variants, genes, and genomic elements that were not studied directly by IGVF. Other interesting public datasets were included as well, including QTLs [18], GWAS results [19], gene annotations [4], protein protein interactions [20,21], curated complexes [22], and linkage disequilibrium [23]. These datasets, their sources, and how they were modeled in the IGVF Catalog are shown in **Table 3**. One of the major advantages of the IGVF Catalog, as compared to similar repositories of project- or consortium-based data, is that this biological context allows immediate interrogation and follow up of discovered relationships. It is very common to supply only shallow data about known objects, such as genes, proteins, or variants. Collecting annotation and relationship data allows users of the Catalog to explore the meaning and context behind the predictions and observed data of the IGVF consortium. Gene annotations, including exon–intron structures, transcription start sites, transcripts, and proteins, were obtained from GENCODE v43 for human and M36 for mice. Genes and proteins are cross-referenced with NCBI and UniProt [24]. Variants, including allele frequencies and annotations were imported from the FAVOR database [17,25], filtered to those present in dbSNP155 [26,27]. Variants that were annotated by public data (GWAS, QTLs) or in IGVF results that were not present in dbSNP155 were added (with no frequency data). Linkage disequilibrium for 4 ancestry backgrounds was imported from TopLD [23]. We have enumerated all possible coding variants that change one amino acid into another by one, two, or three nucleotide substitutions within a codon, even though most of these variants are not observed in population datasets. IGVF has a few predictive models that predict the effect of changing any amino acid to any other throughout the human proteome including MutPred2 [28], and these need to be related to all possible (simple) genomic variations that could produce them. Coding variants are stored separately and linked to their corresponding (possible) genomic variants.

**Table 3.**
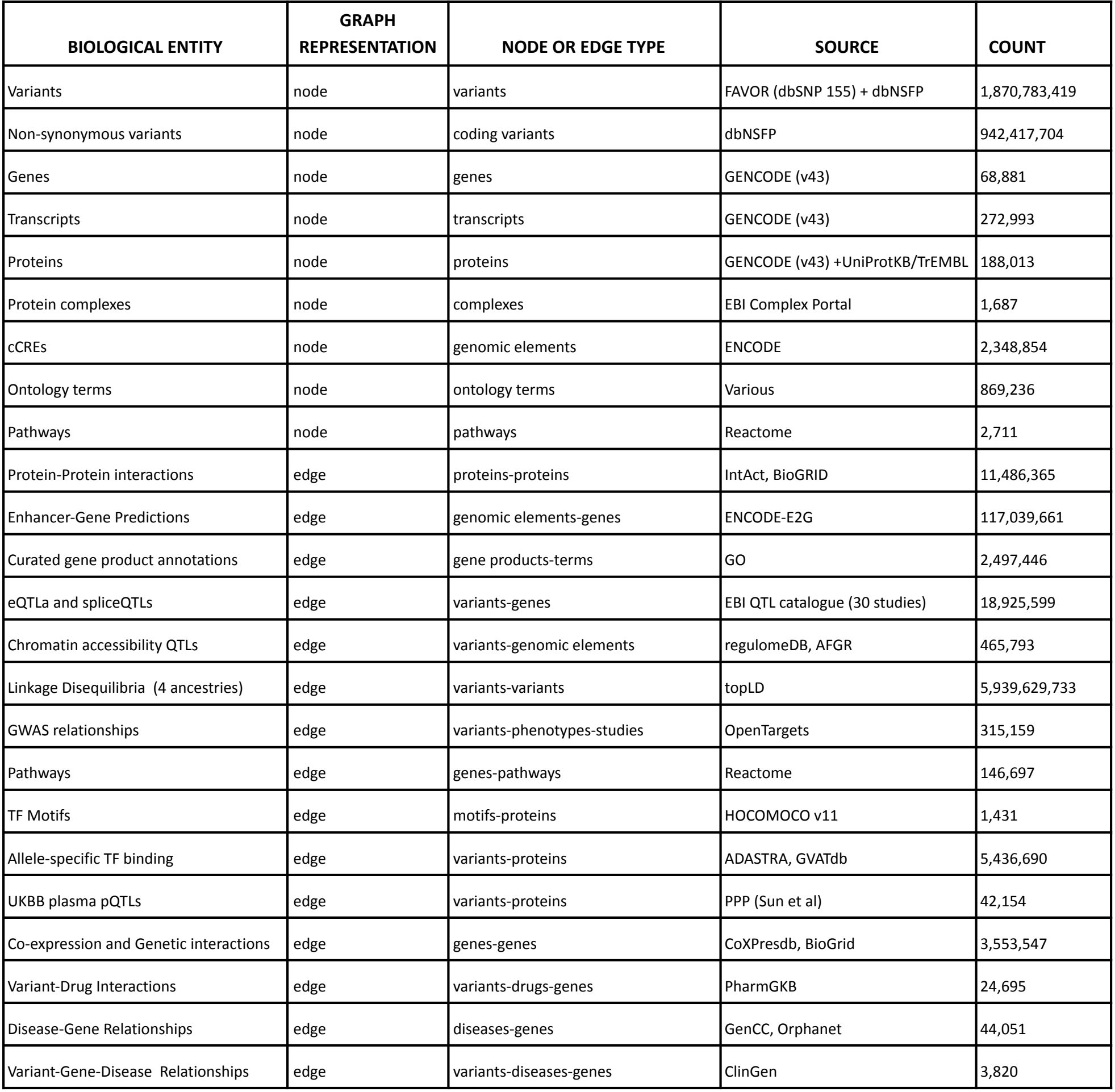
List of Public datasets in the Knowledge Graph. This table shows the number of records loaded from public data sources other than from the IGVF project.

| BIOLOGICAL ENTITY | GRAPH REPRESENTATION | NODE OR EDGE TYPE | SOURCE | COUNT |
|---|---|---|---|---|
| Variants | node | variants | FAVOR (dbSNP 155) + dbNSFP | 1,870,783,419 |
| Non-synonymous variants | node | coding variants | dbNSFP | 942,417,704 |
| Genes | node | genes | GENCODE (v43) | 68,881 |
| Transcripts | node | transcripts | GENCODE (v43) | 272,993 |
| Proteins | node | proteins | GENCODE (v43) +UniProtKB/TrEMBL | 188,013 |
| Protein complexes | node | complexes | EBI Complex Portal | 1,687 |
| cCREs | node | genomic elements | ENCODE | 2,348,854 |
| Ontology terms | node | ontology terms | Various | 869,236 |
| Pathways | node | pathways | Reactome | 2,711 |
| Protein-Protein interactions | edge | proteins-proteins | IntAct, BioGRID | 11,486,365 |
| Enhancer-Gene Predictions | edge | genomic elements-genes | ENCODE-E2G | 117,039,661 |
| Curated gene product annotations | edge | gene products-terms | GO | 2,497,446 |
| eQTLa and spliceQTLs | edge | variants-genes | EBI QTL catalogue (30 studies) | 18,925,599 |
| Chromatin accessibility QTLs | edge | variants-genomic elements | regulomeDB, AFGR | 465,793 |
| Linkage Disequilibria (4 ancestries) | edge | variants-variants | topLD | 5,939,629,733 |
| GWAS relationships | edge | variants-phenotypes-studies | OpenTargets | 315,159 |
| Pathways | edge | genes-pathways | Reactome | 146,697 |
| TF Motifs | edge | motifs-proteins | HOCOMOCO v11 | 1,431 |
| Allele-specific TF binding | edge | variants-proteins | ADASTRA, GVATdb | 5,436,690 |
| UKBB plasma pQTLs | edge | variants-proteins | PPP (Sun et al) | 42,154 |
| Co-expression and Genetic interactions | edge | genes-genes | CoXPresdb, BioGrid | 3,553,547 |
| Variant-Drug Interactions | edge | variants-drugs-genes | PharmGKB | 24,695 |
| Disease-Gene Relationships | edge | diseases-genes | GenCC, Orphanet | 44,051 |
| Variant-Gene-Disease Relationships | edge | variants-diseases-genes | ClinGen | 3,820 |

### API

One of the primary ways of interacting with the IGVF Catalog is via its API (https://api.catalogkg.igvf.org/), and its OpenAPI specification. The API serves as the backend for the Graphical User Interface for the Catalog (https://catalog.igvf.org) but can be used programmatically by remote users and AI agents. Descriptive help for the API is linked on the Catalog site: https://docs.catalog.igvf.org/api-reference/introduction. As an organizing principle, we have added a metadata table called “files_filesets”. This table, or “collection” in ArangoDB parlance details each dataset and file that has been imported into the catalog. Specifically, it links “files” using IGVF (or ENCODE) Data Portal identifiers and a subset of the extensive metadata available on these portals. This collection is organized by “*source*” (IGVF, ENCODE, or one of several public databases), “*lab*” (the IGVF or ENCODE lab producing the data, otherwise “community”), “*class*” (prediction, observed data, or biological relationship), and “*method”*. Method is either the name of the experimental technique used (e.g., “Perturb-seq”), the primary software used in a prediction (e.g., SEMVAR), or an external database name (e.g., ADASTRA [29]). A cross reference to this collection is present in data records as the property `files_filesets` across the knowledge graph. Method, class, source and file_fileset are all filters that can be applied to API endpoints. The API currently has 86 endpoints; 13 for nodes, 20 for edges with IGVF data, 25 “bespoke” (special case, denormalization, or performance optimized) endpoints, 23 for edges with biological context, and 4 utility endpoints, including /files-filesets, the metadata endpoint. Edge endpoints are generally paired; that is there is a “/variants/phenotypes/” endpoint where a variant (or set of variants) is selected and edges to each phenotypes are returned, and a mirror endpoint “/phenotypes/variants/” where a phenotype (including traits and diseases) is selected and edges to each variants are returned. Most endpoints have a “verbose” option which will return selected properties of either or both nodes. Endpoints generally give several consistent ways of selecting input (typically for genes, variants, or elements). Genes and proteins can be selected by their Ensembl ID, gene symbol (name), or alias. Variants can be specified by SPDI [26], HGVS, rsID (not unique, as one rsID can cover multiple alleles), or ClinGen allele registry number [30]. There are both python and R clients (see software availability), along with an Model Context Protocol (MCP) server for agentic AI assistance. Nodes (and edges that link to them) can also be searched by genomic coordinate overlap, or “region”. A region query will take a genomic interval and find all the (genes, variants, elements) that overlap that query and use that to recover matching nodes or edges.

### Implementation Details

The knowledge graph is stored in an ArangoDB (https://arango.ai) graph database. There are essentially 3 components of the software, the database server itself, the API server, and the adapter/loader code. The majority of the code is adapters which interface with the IGVF and ENCODE Data portals, download specific files, parse them and convert them to a series of JSONL files (JSON with added linebreaks), one data record per line. These JSONLs are stored in AWS and can be recreated as needed. Currently we push a new database and API build to production quarterly, while the development version is updated continuously. We maintain a schema (using JSON-SCHEMA) for every node and edge, used to validate every record upon creation. The JSONL files are then ingested into ArangoDB to form the knowledge graph. Future development may use these JSONL files for other backends or database formulations. The API is implemented in TypeScript using tRPC (https://trpc.io/), with an OpenAPI (https://www.openapis.org) specification and Swagger UI (https://swagger.io) supporting interactive exploration and testing.

The codebase is developed with standard software engineering practices. Every change goes through continuous integration on CircleCI (https://circleci.com/), which runs code-style checks, unit tests for the adapters and the API, and integration tests that run API queries against a live development database. Because every adapter’s output is validated against its JSON schema, structural errors in the data are caught before loading as well. The production infrastructure is defined as code using the AWS Cloud Development Kit (CDK, https://aws.amazon.com/cdk/), and the database server machine images are built with Packer (https://developer.hashicorp.com/packer). This means the database cluster and API servers can be deployed automatically and repeatably from version-controlled definitions rather than configured by hand. It also lets us deploy each quarterly release consistently and run development and production environments side by side.

### LLM and MCP integration

The knowledge graph formulation of this database naturally lends itself to creation of an Retrieval Augmented Generation (RAG) for plain text querying. In a RAG system, the user provides an English prompt asking a question about the data in the database. First, the KG schema and the prompt are passed to an LLM (large language model) to create a query in ArangoDB’s proprietary graph language, AQL. This query is then run on the ArangoDB database, and the results are again passed through the LLM to create a reply in English. One of the primary advantages of the RAG method is to constrain the response of the LLM and prevent hallucination. It was straightforward to implement a RAG query system, but limited resources - both in the cost per token for the LLM and both the pressure on the database and amount of data that can be returned has forced us to leave this feature in an experimental mode. We have had better success using agent-moderated LLM queries via Codex or Claude Code. To streamline and moderate this process we created an IGVF Catalog MCP server. The MCP server can be easily added to a coding agent and acts as an interface between the LLM and the Catalog API, so users do not need a deep understanding of either the KG structure or the API. Clarifying questions of the catalog and its capabilities via the API can be expanded upon by the agents themselves as they use various MCP servers, API code bases, and the code base of the catalog itself.

### Case studies

To demonstrate how the IGVF Catalog can help interpret disease-associated genomic regions, here we present three case studies for type 2 diabetes, liver traits, and coronary artery disease. Each case shows a way to integrate evidence from multiple perturbation experiments and prediction methods in the database to prioritize candidate causal variants and propose potential mechanisms. While these case studies do not necessarily represent novel findings, they do confirm previous disease-variant related work and extend specific results to other related findings using the power of the knowledge graph.

#### A FOXA-dependent enhancer variant at the CDC123/CAMK1D type 2 diabetes locus

At the *CDC123/CAMK1D* type 2 diabetes locus, we started from a 500-bp element (chr10:12265775–12266275) tested by CRISPRi in K562 cells from ENCODE [31], which overlaps a distal enhancer-like candidate cis-regulatory element (cCRE) [32] (**Figure 2**). To identify which variant might be functional within the region, we examined the four variants tested by MPRA in IGVF [33]. Only rs11257655 significantly changed reporter activity, where the T allele increased enhancer activity relative to the C allele (log2FC = 0.61) in HepG2 cells. This variant has the strongest association with type 2 diabetes at the locus (p = 10^−63^) [19], with a liver cV2F score of 0.93 [[34]]. It is also a chromatin accessibility QTL (caQTL) in liver [35], consistent with allele-dependent regulatory activity at the enhancer. Transcription factor binding evidence in the IGVF catalog further indicates a potential mechanism for the variant effect on enhancer activity. ADASTRA shows allele-specific binding of the pioneer factor FOXA1 favoring the T allele from ChIP-seq data [29], while SEMVAR also predicts the T allele creates a FOXA1 binding site [36].

**Figure 2.**
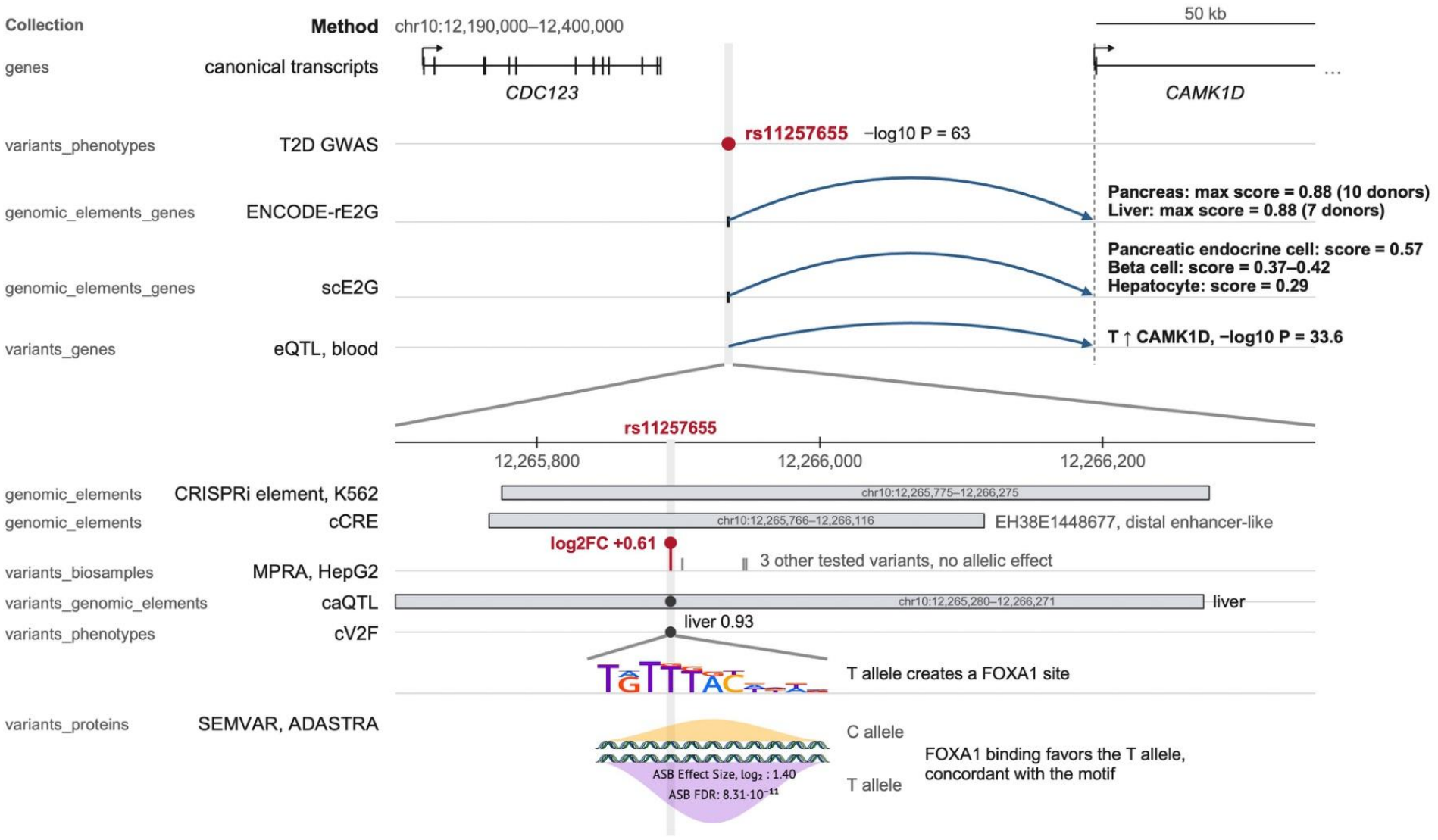
The data in the IGVF Catalog link the type 2 diabetes variant rs11257655 to a FOXA-dependent enhancer of *CAMK1D*. Each track is labelled with its collection name in the IGVF Catalog (left) and method (right). The grey column highlights rs11257655 (GRCh38). **Top:** Arcs show element-to-gene predictions (ENCODE-rE2G, scE2G) and a blood eQTL linking the enhancer to *CAMK1D*. **Bottom:** Bars show the CRISPRi-tested element, the candidate cis-regulatory element (cCRE) and the accessible element in which chromatin accessibility is associated with rs11257655 in liver (caQTL). The T allele increases reporter activity in HepG2 cells, creates a FOXA1 motif and is favored in allele-specific FOXA binding. The motif and allele-specific binding view is adapted from the ADASTRA website (https://adastra.autosome.org/).

We next asked which gene the enhancer regulates. Enhancer-to-gene (E2G) predictions link this enhancer to *CAMK1D*, both from ENCODE bulk data in pancreas and liver tissues [37], and from IGVF single-cell data in pancreatic endocrine and beta cells [17]. In addition, rs11257655 is an eQTL for *CAMK1D* in blood [18], with the T allele being associated with higher expression level. Together, these data in the IGVF Catalog support a mechanism where rs11257655 creates a FOXA binding site in an enhancer, increases *CAMK1D* expression and contributes to type 2 diabetes risk, which was proposed by Fogarty et al. (2014) using independent assays [38].

#### Coding and regulatory variants at liver-trait loci

For liver traits, we started by examining two IGVF CRISPR screens in HepG2 cells that targeted variants associated with HDL and LDL cholesterol [39,40] **(Figure 3)**. In both assays, variant effects were measured through FACS sorting cells based on LDL-C uptake. Specifically, one screen assay used CRISPR interference (CRISPRi) [41] to silence the regulatory region around each target variant, while the other used an adenine base editor which can alter the amino acid sequence [42].

**Figure 3.**
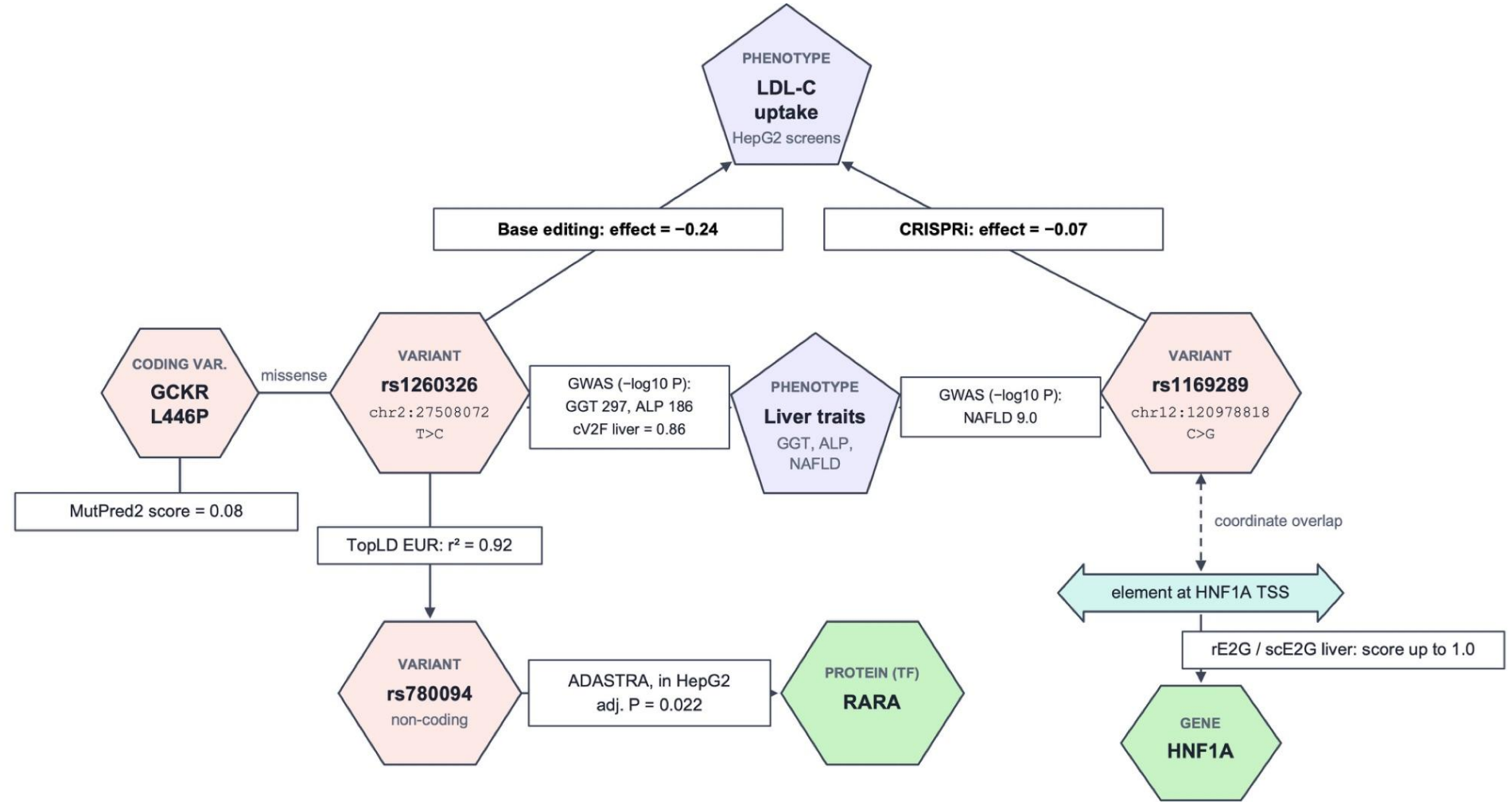
Complementary CRISPR screens reveal coding and regulatory variants at liver-trait loci. Nodes in the graph represent records in the IGVF Catalog node collections, and arrows represent edges between them, with boxes showing the method and data point for each edge. GWAS association is shown as −log_10_(*P*). In HepG2 LDL-C uptake screens, the *GCKR* missense variant rs1260326 (p.L446P) changed uptake from base editing but not from CRISPRi. It is associated with serum gamma-glutamyl transferase (GGT) and alkaline phosphatase (ALP) levels, and its LD proxy variant rs780094 shows allele-specific RARA binding in HepG2 cells. The non-coding variant rs1169289 changed uptake from CRISPRi is associated with non-alcoholic fatty liver disease (NAFLD). It is located in an element at the HNF1A transcription start site, and is linked to *HNF1A* in liver from ENCODE-rE2G and scE2G predictions.

By intersecting significant variants from both assays with liver-trait GWAS variants in the database, we identified rs1260326, which is a missense variant in *GCKR* (p.L446P commonly reported as P446L [43]) as the strongest hit. This variant is associated with serum gamma-glutamyl transferase and alkaline phosphatase, and changed LDL-C uptake from base editing but not from CRISPRi. This is consistent with a coding effect through protein instead of a regulatory effect through local chromatin. Although MutPred2 [28] predicts a modest impact, this variant has a high liver cV2F score [15,28]. Expanding to its LD block, rs780094 (r_2_ = 0.92) shows allele-specific binding of RARA in HepG2 cells from ADASTRA, suggesting an additional regulatory mechanism at the *GCKR* locus.

In contrast, the second hit variant rs1169289 changed LDL-C uptake under CRISPRi (effect = 0.07). It is located at the *HNF1A* transcription start site, which overlaps a genomic element that E2G prediction links to *HNF1A* in liver cells, indicating a regulatory effect on *HNF1A* expression. Overall, this case presents a way to reveal both coding and regulatory mechanisms from complementary assays in the IGVF Catalog.

#### Coronary artery disease: CDKN2A/CDKN2B and NOS3

CRISPRi screens were also performed in two vascular cell types relevant to coronary artery diseases in IGVF (**Figure 4**). In HCASMC-hTERT cells, CRISPRi with TAP-seq [44] measured the effects of 10 candidate cis-regulatory elements on the expression of 7 genes at coronary artery disease risk loci [45]. In TeloHAEC cells, CRISPRi with FlowFISH [31] measured the effects of perturbing a candidate regulatory element on *NOS3* expression [46–49]. By overlapping the elements that significantly changed gene expression in these screens with GWAS variants, we identified the two disease-associated loci with the strongest associations.

**Figure 4.**
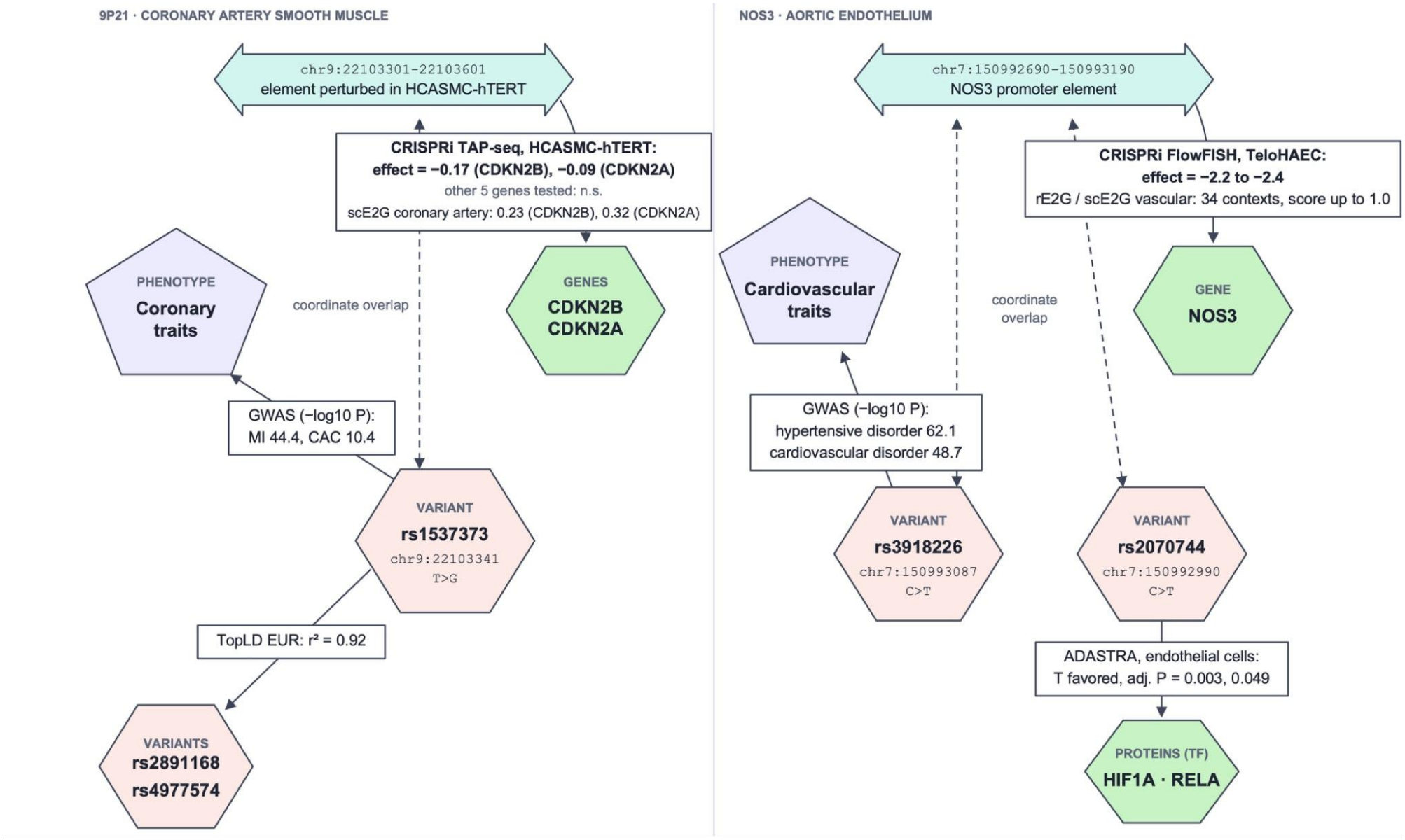
CRISPRi screens in vascular cells connect coronary artery disease variants to target genes. Nodes in the graph represent records in the IGVF Catalog node collections, and arrows represent edges between them, with boxes showing the method and data point for each edge. GWAS association is shown as −log_10_(*P*). In HCASMC-hTERT cells, CRISPRi of an element containing rs1537373 reduced *CDKN2B* and *CDKN2A* expression (TAP-seq). This variant is associated with myocardial infarction (MI) and coronary artery calcification (CAC), and is in LD with the lead CAD variants rs2891168 and rs4977574. In TeloHAEC cells, CRISPRi of the *NOS3* promoter element reduces *NOS3* expression (FlowFISH). Within this element, rs3918226 is associated with hypertensive and cardiovascular disorders, and rs2070744 shows allele-specific binding of HIF1A and RELA favoring the T allele in endothelial cells.

In HCASMC-hTERT cells, CRISPRi of an element containing rs1537373, a variant associated with myocardial infarction and coronary artery calcification, reduced *CDKN2B* and *CDKN2A* expression without significantly changing the other five genes tested. rs1537373 is located in an LD block (r_2_ ≥ 0.8) that includes the lead coronary artery disease variants rs2891168 and rs4977574. E2G predictions in vascular cells link this element to both genes, including single-cell predictions from scE2G in the coronary artery [17].

In TeloHAEC cells, CRISPRi of the *NOS3* promoter element (chr7:150992690-150993190) containing rs3918226 significantly reduced NOS3 expression. This variant is associated with hypertensive disorder and cardiovascular disorder, and E2G predictions link its element to *NOS3* across more than 30 endothelial and vascular contexts. Another variant in this element, rs2070744, shows allele-specific binding of HIF1A and RELA in endothelial cell lines, both favoring the T allele. Together, these examples show a way to combine perturbation screens in disease-relevant cell types with genetic association and regulatory evidence in the database to connect coronary artery disease variants to their target genes.

### Future Directions

The IGVF Catalog Knowledge Graph is a growing and expanding entity. While this release covers the majority of the Characterization data produced by the consortium, experiments and predictions, and new analyses are still being generated. In addition, new and updated releases of public datasets are published continually. There are several directions that we plan to pursue from a data perspective. Specifically, Moore et al have expanded their annotations of ENCODE cCREs [50] to include silencer elements and other biochemical activities; Splice2AI and SpliceVarDB [51] would expand the reach of the catalog into alternative splice forms and their regulation; MorPhic [52] and GREGoR [53] have annotations of disease genes and variants that will be included in later releases. Future work could also include specific gene expression results on a per-tissue and per-cell type basis; These were excluded from the initial releases of the Catalog due to difficult normalization issues. Other data model extensions could include haplotypes, graph genomes, and 3D data from HiC or other approaches. From a software and analysis perspective, building agent driven access to the database to direct questions for visualization and statistics, based on locally trained LLM models will turn the knowledge graph into a fully functional analysis workbench.

## Data and Software Availability

Data files loaded in the latest version of the Catalog are accessible from the IGVF Data Portal with the following query: https://data.igvf.org/search/?type=File&collections=IGVF_catalog_v1.3. All other code for the project can be found on GitHub. Code to reproduce the case studies using the IGVF Catalog API is available as a Jupyter notebook (see **Table 4** for list of software repositories).

**Table 4.** IGVF Catalog Codebase. The GitHub repositories for data loading and the API, MCP server, python and R clients, and the LLM/RAG client.

| **Feature** | **Repository** |
|---|---|
| API and Data loading | https://github.com/IGVF-DACC/igvf-catalog/tree/dev |
| MCP | https://github.com/IGVF-DACC/igvf-catalog-mcp |
| Catalog python client | https://github.com/IGVF-DACC/igvf-catalog-python-client |
| Catalog async python client | https://github.com/IGVF-DACC/igvf-catalog-async-python-client |
| Catalog R client | https://github.com/IGVF/rigvf |
| LLM/RAG flask client | https://github.com/IGVF-DACC/igvf-catalog-llm |
| Case Studies | https://github.com/IGVF-DACC/igvf-catalog/tree/main/case_studies |

## Acknowledgements

This work was supported by the NIH NHGRI IGVF Program (U24HG012012). We thank all the IGVF Consortium members and institutions who have contributed to their experimental and predictive studies and J.M. Cherry, Professor Emeritus.

## Author Contributions

S.D. and I.W. modeled and loaded data into the catalog, developed the API, wrote tex, tablest and figures for the manuscript and defined the case studies, and provided QA support. O.A.J., P.A. and M.L. built, designed, and maintain the catalog knowledge graph software, including API, clients, MCP and LLM/RAG interface J.J., Z. S, KZN, MK, and IG provided data wrangling support for the IGVF Portal K.G. and F.T. built and maintained the IGVF Portal J.M.E., S.L., D.L., MIL are key members of the IGVF Catalog Working group and participated in design, QA, and support of the catalog data base and user interface MIL wrote the Catalog R client, rigvf. T.W. and L.S. provided management support. B.C.H. designed and conceived the catalog knowledge graph, wrote text and edited the manuscript, and manages the Stanford DAC team

---

**Supplementary Table 1 (attached separately)**

Files loaded in the IGVF Catalog (version 1.3). Each row is one entry from the files_filesets collection. The columns show the file and file set accessions, lab, assay, method, class, sample and donor metadata, source and the matching IGVF accessions for ENCODE files.

